# Using the NoiSee workflow to measure signal-to-noise ratios of confocal microscopes

**DOI:** 10.1101/291500

**Authors:** Alexia Ferrand, Kai D. Schleicher, Nikolaus Ehrenfeuchter, Wolf Heusermann, Oliver Biehlmaier

**Author notes:** These authors contributed equally to this work (Alexia Ferrand, Kai D. Schleicher).

## Abstract

Confocal microscopy is used today on a daily basis in life science labs. This “routine” technique contributes to the progress of scientific projects across many fields by revealing structural details and molecular localization, but researchers need to be aware that detection efficiency and emission light path performance is of major influence in the confocal image quality. By design, a large portion of the signal is discarded in confocal imaging, leading to a decreased signal-to-noise ratio (SNR) which in turn limits resolution. A well-aligned system and high performance detectors are needed in order to generate an image of best quality. However, a convenient method to address system status and performance on the emission side is still lacking. Here, we present a complete method to assess microscope and emission light path performance in terms of SNR, with a comprehensive protocol alongside NoiSee, an easy-to-use macro for Fiji (available via the corresponding update site). We used this method to compare several confocal systems in our facility on biological samples under typical imaging conditions. Our method reveals differences in microscope performance and highlights the various detector types used (multialkali photomultiplier tube (PMT), gallium arsenide phosphide (GaAsP) PMT, and Hybrid detector). Altogether, our method will provide useful information to research groups and facilities to diagnose their confocal microscopes.

## Introduction

Confocal microscopy of fluorescently labelled specimen has become an increasingly used and important tool in biological research across disciplines. Proper results rely on accurately set and aligned microscope systems, images of which are often evaluated for high-resolution structural information but also intensity content. While in a perfect optical system the resolution is in theory only limited by the objective numerical aperture and wavelength used, the practical resolution limit is reached when the specimen signal is indistinguishable from the instrument noise^1^. It has been standardly accepted to calculate resolution by using bright and well separated fluorescent point sources to measure full width half maximum (FWHM), whereas a more direct alternative to measure resolution would assess the distance of two rather dim fluorescent point sources (Rayleigh criterion,^2,3^). For accurate results, it is hence imperative to have a signal that is well distinguishable from noise, i.e. a high signal-to-noise ratio (SNR). Maintaining a well-adjusted system by monitoring the SNR is an important step that can give valuable information about the quality of the system, its proper alignment, its sensitivity and the overall system status. Therefore, SNR is a key factor when a researcher is choosing a microscope to work with, which becomes especially relevant in a facility environment where several systems of different age and vendor may be present. Assessing SNR as part of a general monitoring routine together with measurements of laser intensity and point spread functions (PSF) is therefore important but has been a tedious task so far, as previously described methods to address SNR lack ease of use^4,5^. Some useful tools such as ConfocalCheck help to monitor confocal performances^6^. But whereas the whole purpose is globally the same, ConfocalCheck gives results spanning from laser stability, objective chromatic aberrations, to galvo stability, but does not address emission light path performance and SNR.

A central element of the emission light path contributing to SNR is the detector used in a given setup. In this paper, we have tested systems including three different types of detectors, namely the classical photomultiplier tubes (PMT) involving photosensitive elements (photocathodes) made from antimony-sodium-potassium-caesium (known as multialkali PMT, S-20) or gallium arsenide phosphide (GaAsP PMT), and the more recent hybrid detectors (HyD). While multialkali PMTs have been the standard in confocal microscopy for a long time, more recent materials like GaAsP have superior quantum efficiencies (QE) in the visible spectrum and represent the latest generation of photocathodes used by vendors^7^.

Photons emitted by a fluorescent sample for example hit the photocathode, thereby releasing electrons (called photoelectrons) from the cathode in a process known as the photoelectric effect. Due to the quantum nature of light, the number of photons arriving at the photocathode in a given time interval is subject to statistical fluctuations described by a Poisson distribution. The uncertainty of this distribution (i.e. noise) is known in this context as photon shot noise and represents the fundamental limit of the SNR. The efficiency of converting an incident photon to a photoelectron is described by the QE of the photocathode material, i.e. the ratio of photoelectrons to incident photons^8^. However, a single photoelectron is difficult to measure and hence requires amplification by the detector in order to produce a definite output.

In PMTs, amplification of each photoelectron is achieved via a series of dynodes. The magnitude of this amplification can be controlled by applying a voltage (often called “gain” in a systems software, ∼800 V across a series of dynodes^7^) to accelerate the photoelectron towards the dynodes which creates multiple secondary electrons upon impact based on their kinetic energy. While this process leads to a greatly enhanced signal, the multi-stage amplification at several dynodes introduces an uncertainty in the height of the output pulse as the number of secondary electrons created at each dynode is not constant but also follows a Poisson distribution. This statistical fluctuation in the generation of secondary electrons is known as multiplication noise.

To overcome this drawback, novel hybrid detectors combine GaAsP-based photocathodes for light conversion followed by a single-step acceleration with high voltage (∼8 kV). Similar to the amplification process in an avalanche photodiode, the accelerated photoelectrons in a HyD impact a semiconductor material and a subsequent multiplication layer, resulting in an “avalanche” of many secondary electrons. This high-gain single-step amplification greatly reduces the multiplication noise in hybrid detectors.

Besides incident photons, thermally generated electrons at the photocathode as well as at the dynodes also result in an output signal and are indistinguishable from photoelectrons. The signal from thermally generated electrons is referred to as the “dark noise” or “dark current” of a detector and pose a problem especially for weak signals. Due to their smaller sized cathodes^9^ and the absence of dynodes, HyDs have a lower dark current and are therefore well suited for dim samples and even photon-counting applications.

In a detector, noise is the variation in output when given a constant input signal; it is not to be confused with the difference between signal and background in an image. From a microscopist’s point of view, the background may be viewed as an omnipresent offset in signal intensity that equally applies to the whole image, for example stray light entering the detector. From a biologist’s point of view, the background is often referred to as regions in the specimen that show autofluorescence or unspecific staining, for example unspecific binding of primary antibodies or low amounts of fusion proteins expressed in cell compartments or tissues other than the regions of interest. While the prior definition of background is of considerable importance for the final image quality, the latter is not a property of a given microscope and is hence not included in the background as defined here. However, for a given signal to be detectable it needs to be well above the background and hence the signal-to-background ratio (SBR) is another important metric when assessing image quality. It is noteworthy that in the absence of any other factor, i.e. stray light, the background can be viewed as a measure of the dark current.

The main goal of this study is to provide research groups and facilities with the protocols and tools necessary to conveniently assess the quality of their confocal microscope and to be able to compare instruments amongst each other. When doing the measurements, we focused on imaging conditions that are relevant for fixed biological samples, i.e. low laser powers that minimize photobleaching and photodamage. Hence our measurements are not meant to achieve the highest SNR values possible but rather reflect the state of the system when operated under typical working conditions.

## Results

### Acquisition protocol and NoiSee macro

To assess the image quality and SNR of confocal systems, we defined a precise workflow comprised of several steps (Figure 1). To have comparable SNR measurements between instruments, many parameters need to be set. (i) The objectives need to have the same numerical aperture (NA), and the samples need to be illuminated with a fixed photon dose e.g. (ii) with a fixed pixel dwell time and (iii) a fixed laser intensity at the focus, (iv) a fixed pinhole diameter (same back projected pinhole^10^), and (v) a fixed detection range (Figure 1, box). When used for comparing a single system over time, the values of these parameters can be set with more flexibility, but will still have to be kept constant between the measurements.

**Figure 1.**
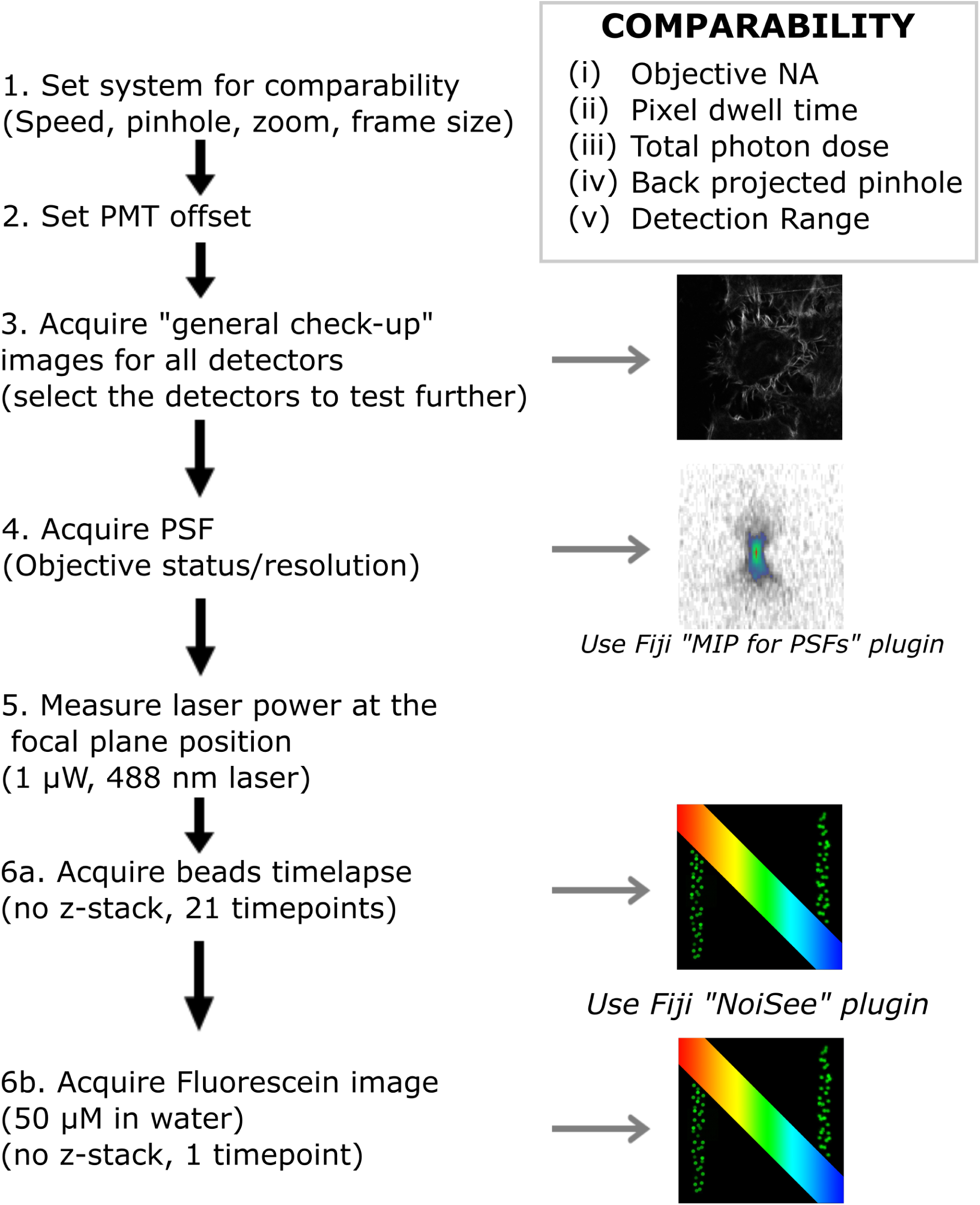
Workflow describing how to set up the confocal for SNR measurements. After carefully setting up the system and detectors (steps 1-2), a “general check-up” image is recorded on all detectors using a well-known standard sample and the resulting image is representative of the image quality of the system (step 3). In step 4, a PSF is recorded on any chosen detector and analysed using the “MIP for PSFs all microscopes” macro in Fiji. The SNR of a given detector is then addressed by setting the laser power in focus to 1 µW (step 5) and alternatively recording a 2D timelapse of beads (step 6a) or a single 2D image of a homogeneous fluorescein solution (step 6b), which can both subsequently be analysed using the “NoiSee” macro in Fiji.

To achieve comparability between systems, we adjusted the pinhole size, the scanner speed, the zoom and frame size in the respective software (Figure 1, step 1). A table with all the settings for comparability can be found in the Supplementary Material (Supplementary table 1). When adjusting the dynamic range of the detector it is important to ensure no clipping of the background/dark current. Therefore, to make sure that there will not be any difference in the measurement of the SBR, we set the offset in a similar way. The PMTs offset was set to the highest possible level avoiding zero-values when the laser is off (Acousto-Optic Tunable Filter (AOTF) and shutter closed) (Figure 1, step 2). For the HyD detectors, no offset adjustment is necessary as no background/dark current was detected.

We imaged HeLa cells stained for actin filaments (Phalloidin-Alexa 488), under conditions that are standardly used for imaging fixed samples on a confocal microscope (“general check-up image”, Figure 1, step 3). The signal was set to be just under saturation, as described in the previous studies^4^. This initial image helps to give a first idea on the detector performance. For the present study, at this step, we selected the detector that was giving the best image.

The objective is crucial for the SNR measurement. To make sure that the objective used is of good quality we checked its PSF (Figure 1, step 4) by using the well-established PSF distiller macro “MIP for PSFs all microscopes” to calculate the PSFs^11^. In case the lens shows a deformed PSF, it should be cleaned further, or sent for repair in case of a more serious defect. Only with a good PSF, meaningful and comparable SNR measurements can be obtained.

The power at the focal plane was set to 1 µW, which on one hand represents a typical power value for imaging a biological samples and on the other hand does not introduce saturation of our 1 µm bead test sample (Figure 1, step 5). To achieve comparable laser power measurements on different systems, we accounted for their specific blanking times.

A full description of how the laser power measurements were set can be found in the Material and Methods section. The SNR/SBR calculations were done using NoiSee, our easy-to-use macro for Fiji^12,13^. It automatically calculates SNR, SBR and provides further quality measures to assess drift and photobleaching. We tested two ways to evaluate the SNR. The first method uses a commercially available TetraSpeck 1 µm fluorescent beads slide (as described in Figure 1 step 6a and below) whereas the second method uses a uniform fluorescein solution (Figure 1, step 6b). In brief, in the fluorescein method, the mean intensity of a single 2D image of such a solution (i.e. signal) alongside its corresponding standard deviation (i.e. noise) provides a quick measure of the SNR when accompanied by an image taken without illumination (dark image). Here every pixel of the final image can be considered as a separate “timepoint”. The mean of the dark image provides a measure of the background. The setup for the fluorescein method and the results from NoiSee are presented in Supplementary Tables 2 and 3 and Supplementary Figure 1.

For simplicity, only values derived from the beads method are reported in the following results. Fields of view (FOV) containing between 25-50 beads were imaged as single plane time-lapse for 21 timepoints. From the line profile of a bead (Figure 2a), the SNR is as the average signal (*μ*_s_) divided by its standard deviation (*σ*_s_), *SNR* = *μ*_s_/*σ*_s_, after background subtraction (Figure 2b,^14,15^). Figure 2b shows how these quantities can be obtained by monitoring the brightness of a fluorescent bead (signal) and variation thereof in time (noise), given there is no bleaching during acquisition.

**Figure 2.**
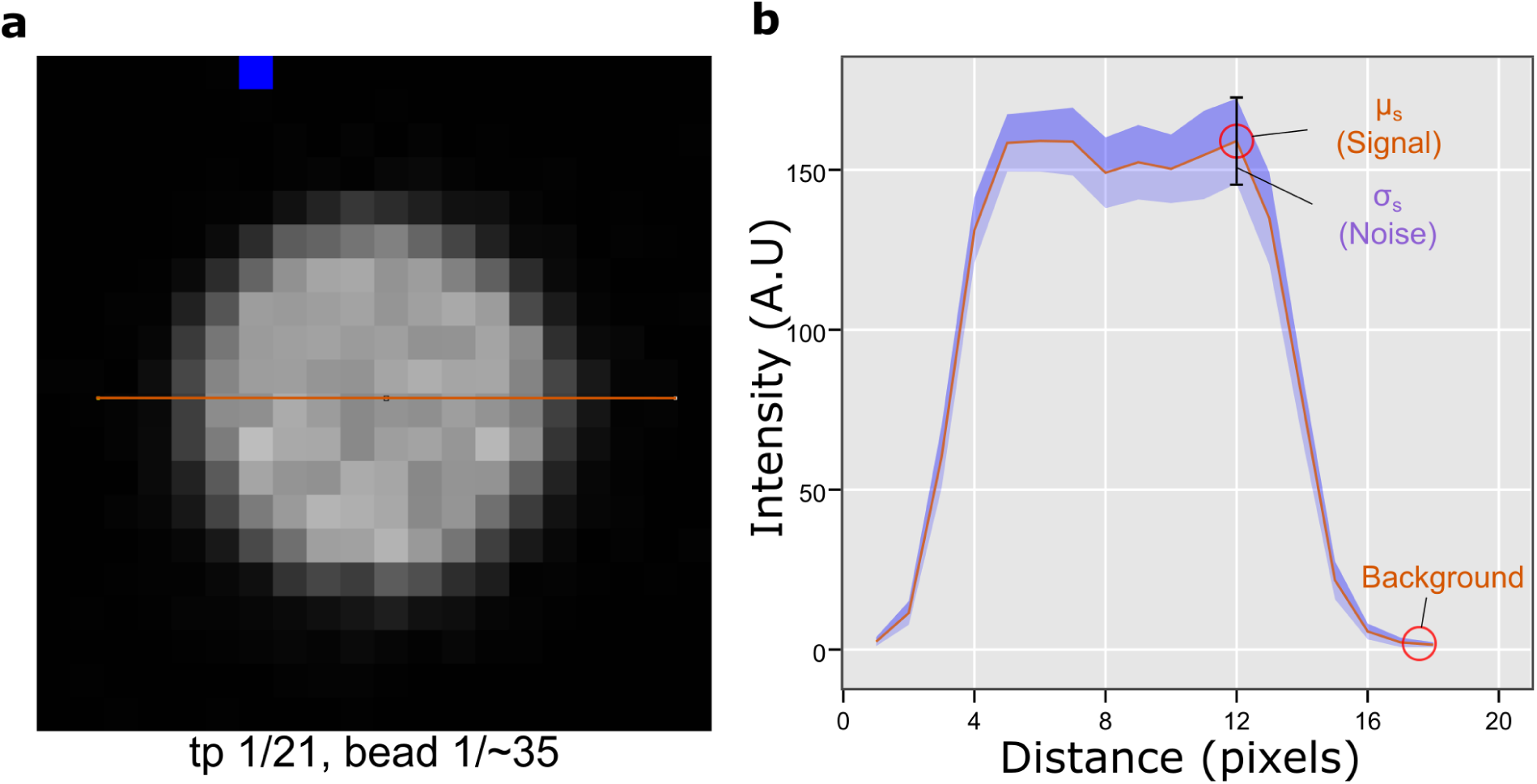
Definition of the signal-to-noise ratio derived from images of fluorescent beads. a. Zoom in on a single bead (timepoint 1) from a sequence of 21 raw images. The orange line indicates the measurement area for the plot presented in b. b. Mean intensity profile (orange) of the bead as shown in a alongside its standard deviation across all timepoints (purple). The peak value of the mean intensity is defined as the “signal”, while its accompanying standard deviation is defined as “noise”. The low intensity area region next to the bead is defined as “background”.

After bead segmentation and subtraction of the average background (Figure 3a), NoiSee calculates the average intensity and associated standard deviation of the timelapse (Figure 3b, c) and subsequently builds their ratio, i.e. an image of the SNR (Figure 3d). From the average intensity image, the brightest pixel per bead is identified and its intensity measured on the SNR image, resulting in several individual data points per image. Bleaching and sample drift are monitored by following the beads mean intensity and associated standard deviation in the raw data image over the course of the entire time lapse and are plotted accordingly. Changes in mean intensities point to bleaching or drift in z (axial). For a visual representation, the macro generates kymographs of selected beads (Figure 3e, f). Resulting values are presented in a summary table and are saved automatically alongside with measurement points and regions from the ROI manager (Figure 3g, h). The latter ones are available for interactive review after the macro has completed. NoiSee additionally offers to save individual data points in text files as well as a PDF summary file including all images and graphs generated (see the NoiSee user guide in the supplementary material).

**Figure 3.**
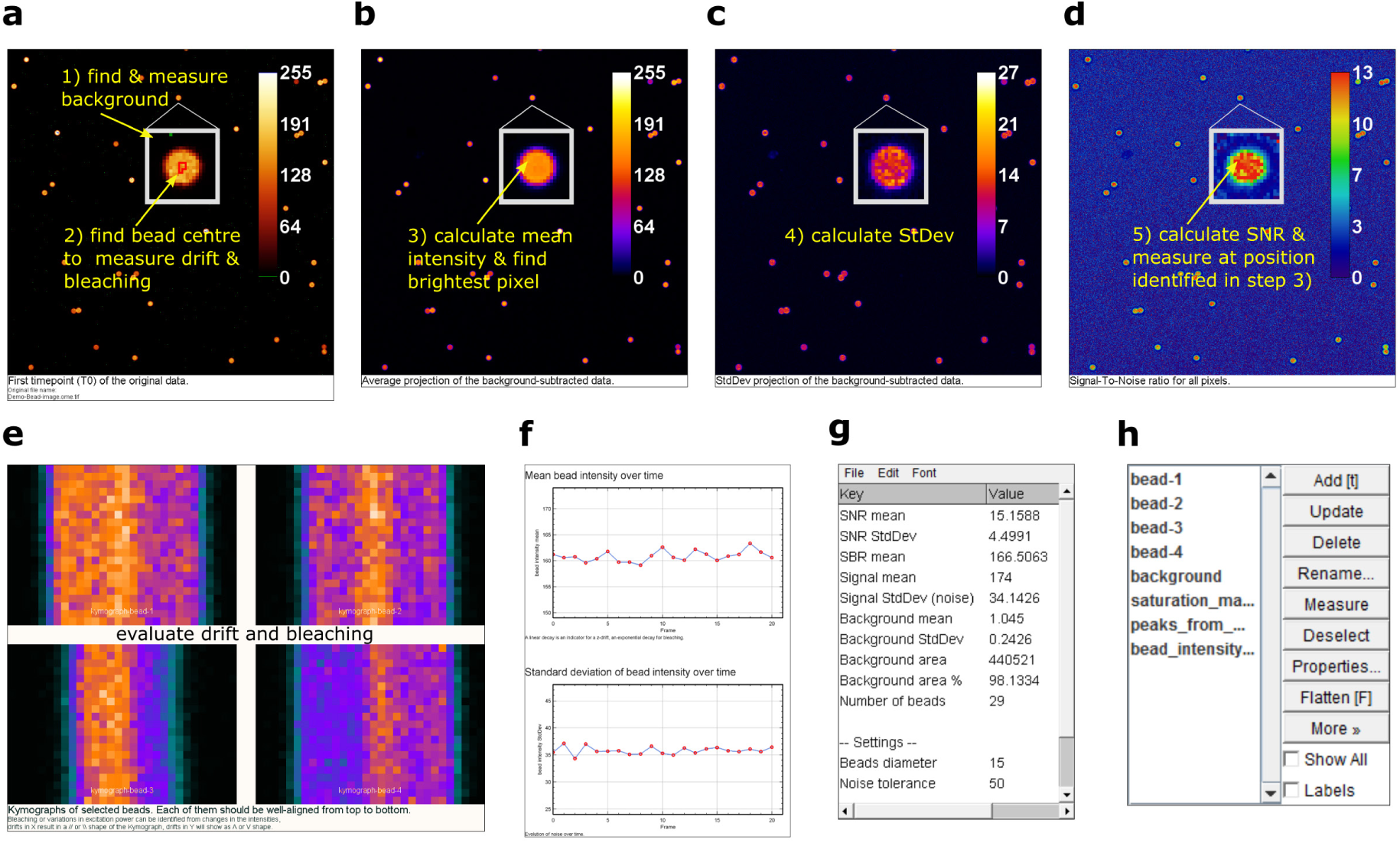
Screenshots of NoiSee output and features. a. NoiSee automatically segments beads, their centre area and the image background in the raw data. b. Average intensity of the raw data projected over time, after background subtraction. Each pixel corresponds to the mean intensity of the raw data at the respective position. NoiSee automatically identifies the brightest pixel per bead on which the SNR is measured. c. Standard deviation of the raw data projected over time, after background subtraction. Each pixel corresponds to the standard deviation in intensity of the raw data at the respective position. d. A visual representation of the signal-to-noise ratio is generated by dividing the average intensity projection from b by the standard deviation projection from c. NoiSee automatically measures the SNR value for each bead at the pixel values identified in b. e. Kymographs corresponding to the cross-section of automatically selected beads to aid visual inspection of lateral sample/stage drift and bleaching. f. Plots of the mean bead intensity and standard deviation over all bead centre areas to evaluate axial drift and bleaching. g. Results table presenting a summary of the calculations. h. ROI manager including all measurement regions for interactive inspection.

Combining this information with the general check-up image and the PSF allows for an “ID-card” of the systems, as displayed in Supplementary Figure 2.

### Example images of high and low SNR

Figure 4 displays the standard images acquired on different microscopes under the same conditions from the step 3 of our method (“general check-up” images). It is important to point out that the standard images presented in Figure 4 are not acquired with exactly the same conditions as the beads from which the NoiSee score is calculated (step 6a). They are representative versions that include two-fold averaging and higher laser power to make sure that the dynamic range of the detector is fully filled, i.e. just below saturation. They match the images that scientists would acquire on the specific machines. The corresponding NoiSee scores were calculated as described in the method (step 6a, at 1 µW without averaging) and the impact of high or low signal-to-noise ratios on image quality is shown. When the image of the highest score (Figure 4f) is compared to the image of the lowest score (Figure 4j), it is obvious that the fine structure details are lost in high noise conditions.

**Figure 4.**
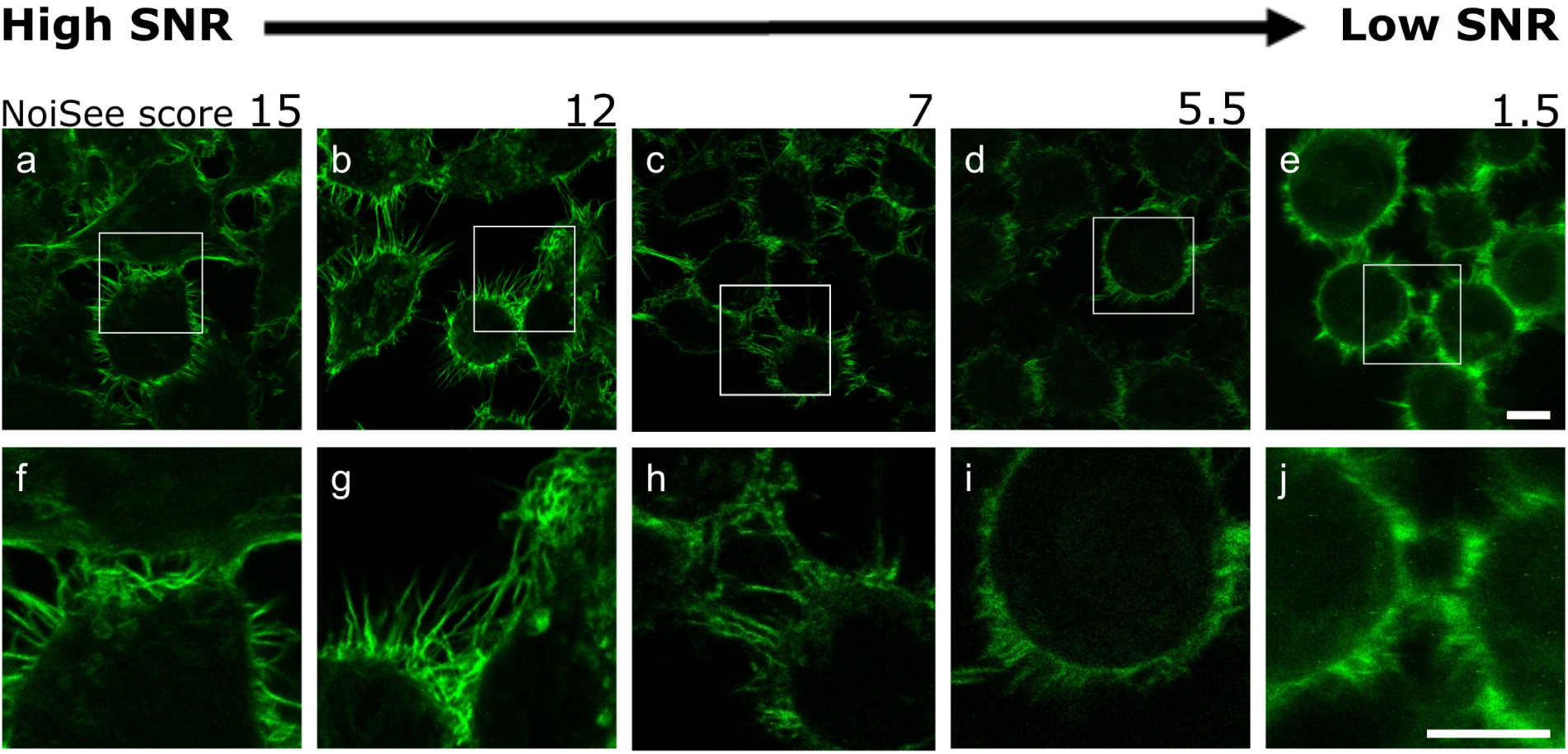
Image quality as a function of SNR. Cells with actin fibres imaged on different confocal systems and showing decreasing SNR scores from left to right. The detectors were chosen to represent the range of SNR score: a/f. LSM800 GaAsP2; b/g. LSM700up PMT2; c/h. SP8M PMT3; d/I. SP5II PMT1; e/j. SP5 MP HyD2. Scale bar 10 µm.

### NoiSee analysis results and pertinence

Table 1 and Figure 5 summarize the results obtained for five CLSM with respect to the detector type used. Firstly, emission light paths using novel GaAsP-based PMTs outperform the ones based on conventional multialkali PMTs in term of their SNR (figure 5a and 5c). Secondly, HyDs-associated emission light path show background that is two orders of magnitude lower than from any PMT-associated emission light path, i.e. virtually zero. Consecutively, systems equipped with HyD detectors score the highest SBR values (figure 5b). Thirdly, comparing Leica detectors amongst each other reveals the superior quality of the hybrid technology. Closer inspection of the recorded noise alone shows that the LSM800 GaAsP-PMT-associated emission light path is performing better in terms of its noise level compared to the LSM700 multialkali PMTs ones (figure 5d). It is however important to note that it is not possible to compare absolute SNR measurements for individual detectors directly, especially between microscopes because of the different amplification technologies and gain scaling between PMTs, HyDs and manufacturers.

**Table 1.**
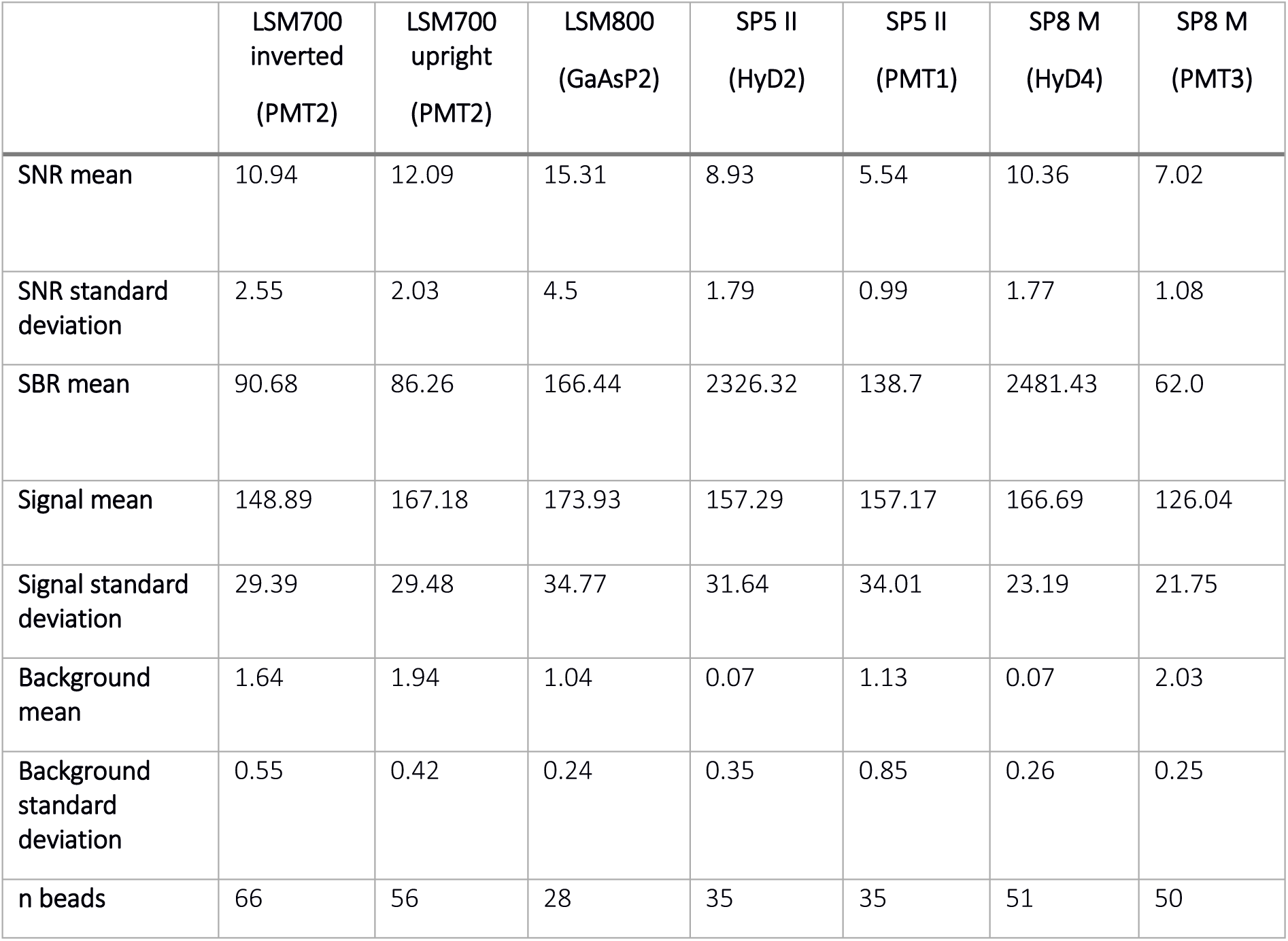
Results at the different microscopes using the Beads method

|  | LSM700<br>inverted<br>(PMT2) | LSM700<br>upright<br>(PMT2) | LSM800<br>(GaAsP2) | SP5 II<br>(HyD2) | SP5 II<br>(PMT1) | SP8 M<br>(HyD4) | SP8 M<br>(PMT3) |
| --- | --- | --- | --- | --- | --- | --- | --- |
| **SNR mean** | 10.94 | 12.09 | 15.31 | 8.93 | 5.54 | 10.36 | 7.02 |
| **SNR standard**<br>**deviation** | 2.55 | 2.03 | 4.5 | 1.79 | 0.99 | 1.77 | 1.08 |
| **SBR mean** | 90.68 | 86.26 | 166.44 | 2326.32 | 138.7 | 2481.43 | 62.0 |
| **Signal mean** | 148.89 | 167.18 | 173.93 | 157.29 | 157.17 | 166.69 | 126.04 |
| **Signal standard**<br>**deviation** | 29.39 | 29.48 | 34.77 | 31.64 | 34.01 | 23.19 | 21.75 |
| **Background**<br>**mean** | 1.64 | 1.94 | 1.04 | 0.07 | 1.13 | 0.07 | 2.03 |
| **Background**<br>**standard**<br>**deviation** | 0.55 | 0.42 | 0.24 | 0.35 | 0.85 | 0.26 | 0.25 |
| **n beads** | 66 | 56 | 28 | 35 | 35 | 51 | 50 |

**Figure 5.**
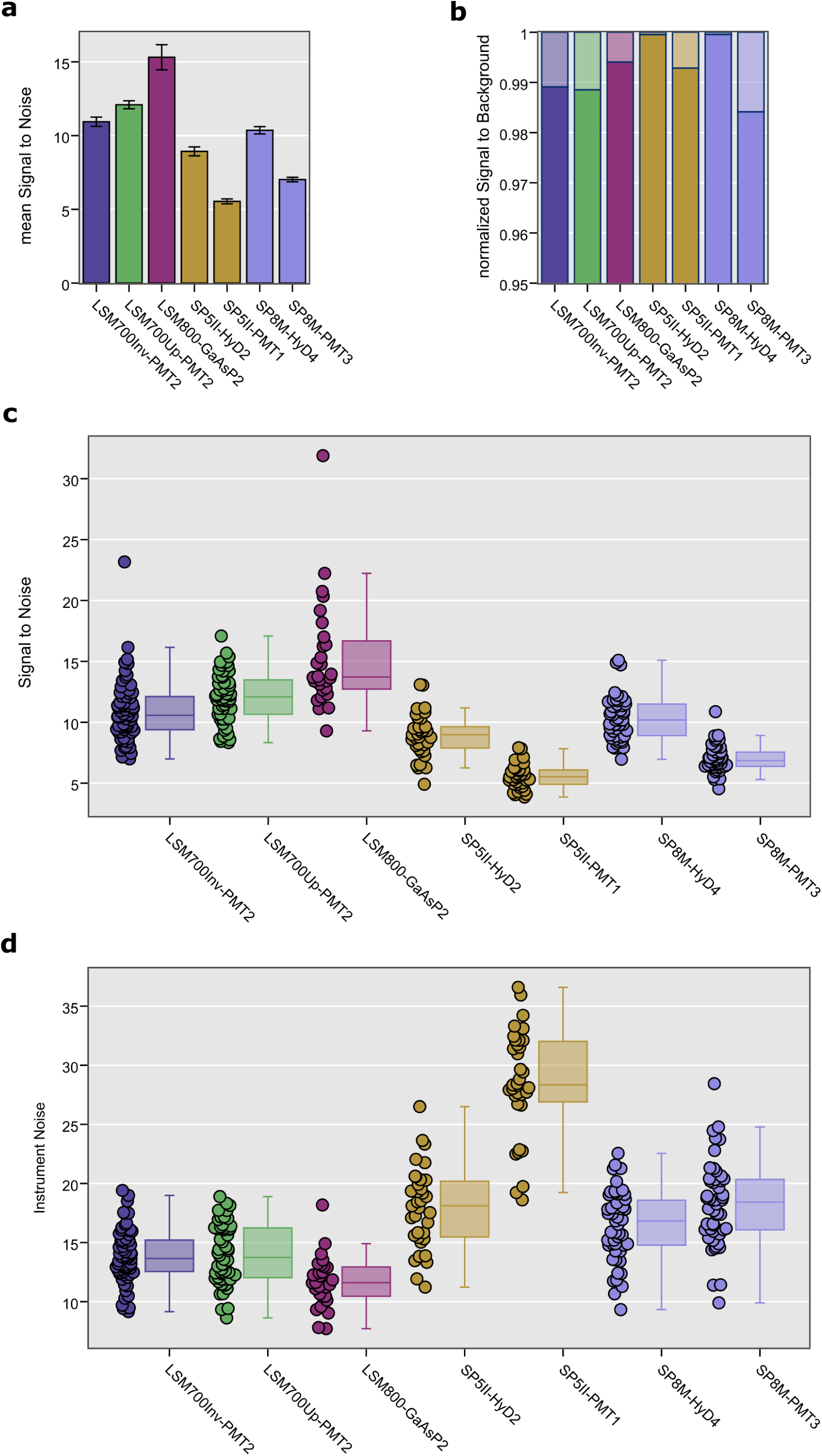
NoiSee results of five different confocal laser scanning microscopes equipped with different detector technologies. a. bar charts of mean SNR reveal significant differences between the seven different detectors tested. Error bars represent the standard error of the mean. b. normalized NoiSee SBR scores of the same systems and detectors. Solid colour denotes the mean signal while transparent colour denotes the mean background. c. box plots summarizing the distribution of all individual SNR values (one per bead) from all beads in an image as measured by NoiSee (solid circles next to the boxes). d. box plots describing the distribution of all noise values that correspond to c. All data points underlying these graphs is a direct output of NoiSee.

To ensure that obtained SNR results are not underestimated due to photobleaching we plotted the average bead intensity per frame (Supplementary figure 3), which revealed no significant decay over time or between measurements and thus ensures the comparability between datasets.

The reproducibility of the measurements was addressed. Figure 6 shows the results for two individual systems (LSM700up/LSM700inv) as measured a few days apart (Repetitions 1 and 2, stated as R1 and R2). Analysis of the NoiSee results reveals a significant difference between the measurements on the LSM700inv (Figure 6a and c) while no difference was found for the LSM700up. Inspection of the instrument noise distribution presented in Figure 6d shows that this difference does not result from a higher noise level and must therefore be attributed to the difference in the average signal recorded (see supplementary table 4). In comparison, SBR values were generally constant, indicating no change in sample background or dark current was present, which constitutes less than 2% of the overall intensity measured in all cases (Figure 6b). In the present instance, we cannot exclude that the beads imaged for R2 were bleached compared to the ones in R1 due to previous usage, or that this result reflects the laser stability of the system, which could explain the slight discrepancy.

**Figure 6.**
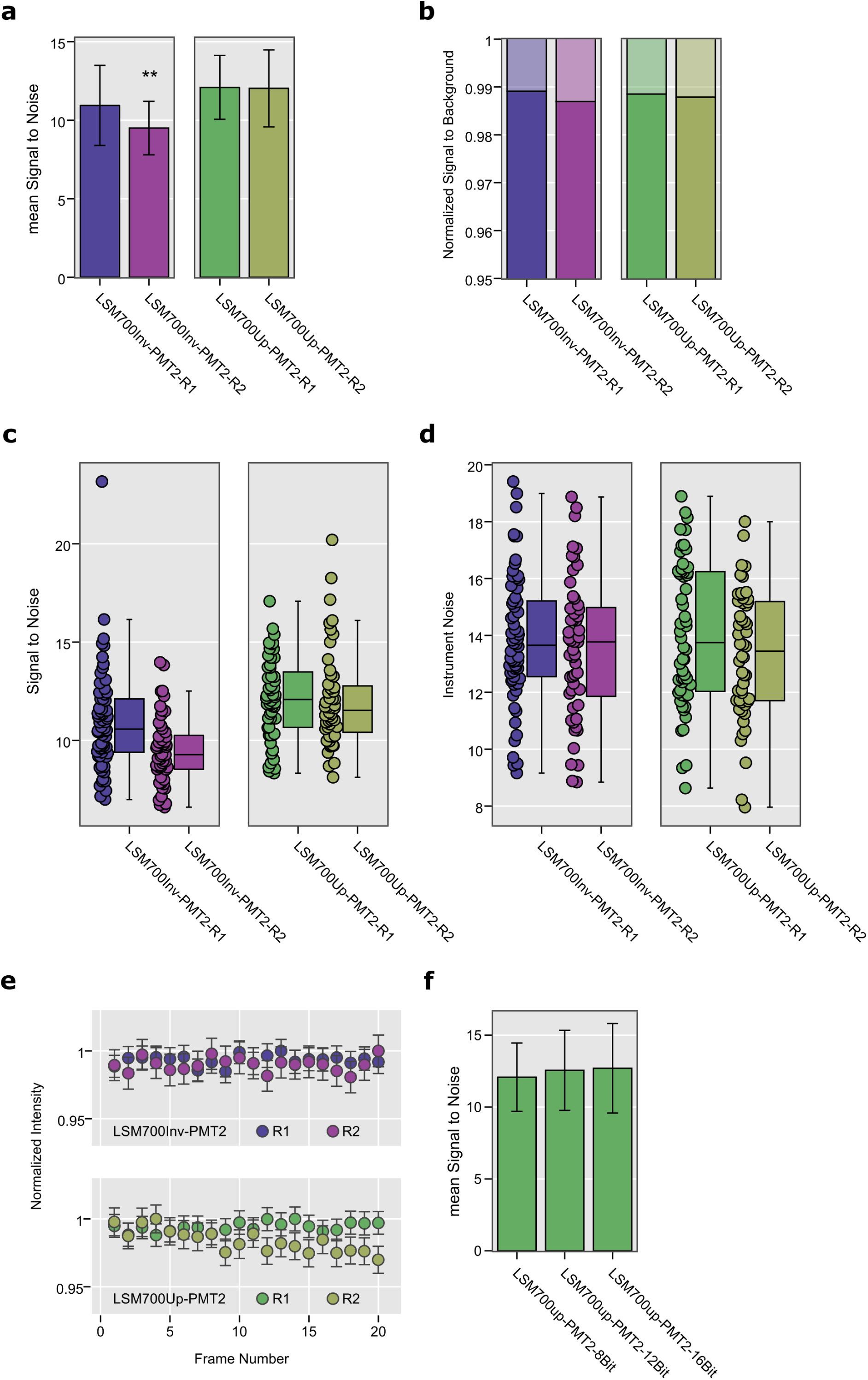
Repeated measurements show the reliability of NoiSee score. a. Bar charts of the mean SNR score of two individual systems (LSM700inv, blue/purple and LSM700up, green/olive) measured with a few days between measurements (Repetitions 1 and 2, R1 and R2). Significance was calculated using Welch’s t-test for independent samples^34^. P-values in a for the LSM700inv were p=0.0004 and p=0.8930 for the LSM700up measurements, respectively. b. SBR of the measurements corresponding to the data presented in a. The distribution of individual values is summarised in the box plots presented in c (SBR) and d (noise). e. Normalized mean intensity across all beads as measured by NoiSee for the two LSM700 systems across the two repetitions. The intensity fluctuations generally stay within the measurement error, proving that beads did not bleach during acquisition, except the slight bleaching in the second repetition of the LSM700up measurement (olive, bottom panel). f. Bar charts of the mean SNR score for the LSM700up recorded at 8-,12-, and 16-bit, respectively. Welch’s t-test for independent samples revealed no significant differences [p(8-bit vs. 12-bit)=0.33, p(8-bit vs. 16-bit)=0.23, p(12-bit vs. 16-bit)=0.79]. Error bars represent the standard deviation (a, f) or the standard error of the mean (e).

We further tested if the bit-depth of the acquired image influences the SNR score. NoiSee accepts 8-, 12-and 16-bit images. To simplify bead detection, NoiSee includes a rescaling step to 8-bit that ensures ease-of-use and consistency of results (see NoiSee user guide in the Supplementary Material). No rescaling step is required for the fluorescein part of the macro. Beads were acquired on the LSM700Up at 8-bit, 12-bit and 16-bit. The offset was adjusted for each bit-depth. The SNR scores were both similar to each other and to the measurement for the same instrument in Figure 6a, with no significant differences between the various bit-depths (Figure 6f, Supplementary Figure 4 and Supplementary Table 6). These results show that the acquisition bit-depth does not influence the SNR score.

## Discussion

In facilities and all laboratories doing high-end microscopy, it is important to assess the performance and image quality of the microscopes. This process includes quality control on individual systems over time as well as inter-system comparisons. Many criteria can/should be routinely tested, such as illumination uniformity across FOV (using a fluorescein solution, or following the protocol described by Brown and coworkers^16^), XYZ chromatic aberrations and resolution (using the PSFj^17^ or MetroloJ^18^ tools, or using 0.2 µm beads as detailed by Cole and colleagues^19,20^), laser power and stability (using power meter/time series). Maintaining a well-adjusted system by monitoring the SNR is an important step that can give valuable information about the quality of the system, its proper alignment and overall emission light path, especially as the SNR is the limiting factor in image quality and ultimately limits resolution. However, despite the importance of this metric, its measurement has proven difficult in the past due to a lack of a straightforward method. Our comprehensive method to assess microscope and detector performance in terms of their SNR can now fill the gap, and provides a helpful tool for life science labs and facilities.

Many different tools have been successfully used in the past to measure SNR for different types of microscopes and flow cytometers^4,5,14,15,21-24^. The method presented here builds on these approaches and additionally offers a ready-to-follow workflow alongside a freely available analysis macro; it can serve as an easy-to-use tool for facilities and high-end confocal microscopy labs to be integrated into their standard routine. Combined with other tools for quality check such as Sectioned Imaging Property (SIP) charts^25,26^ or ConfocalCheck^6^, NoiSee allows for streamlined full assessment of the system status.

While it is rather hard to set all parameters 100% equivalent between different systems, an extreme care was taken to make sure that all parameters were as comparable as they could be. We are confident that the overall scores are representative of the emission path sensitivity of our systems. Our results are not intended to reflect the overall quality of the respective technologies and/or microscope vendors, but rather reflect the state of our microscopes emission light path at a specific point in time. We cannot exclude that the slight variations in back projected pinholes or gain values have a minor effect to the NoiSee scores. We have tested five different microscopes, each equipped with one or more detectors of three different types, namely the classical multialkali PMTs or GaAsP PMTs and the more recent hybrid detectors (HyDs). Our results show that GaAsP PMTs and HyD detectors display a higher SNR than their respective standard PMT counterparts. The higher SNR of the GaAsP-PMTs compared to their multialkali-counterpart is expected and likely reflect the enhanced QE of the novel photocathode materials^7^. Due to their design, HyDs do not suffer from multiplication noise compared to PMTs, which results in a higher SNR score. This becomes especially apparent if both detector types are available in the same microscope. Different SNR between detectors using the same photocathode material may reveal differences in the number of photons delivered to the photocathode (e.g. the light path in general or the alignment thereof), or overall PMT build-up. It is noteworthy that detector design may vary greatly between systems and could hence also show different effective QEs, that is the overall QE of the detector rather than the QE of the photocathode alone^7^. While all detectors tested have a background lower than 2% of the signal, our results confirm that HyD detectors are extremely sensitive and have virtually no background; consecutively they score the highest SBR. The standard deviation of the background is a less intuitive metric as it does not clearly enter the SNR or SBR. Here, a high value may indicate a mechanical or electronic defect on the detection side, such as a pinhole misalignment (data not shown). These results are consistent when checked with an alternative method utilizing an aqueous solution of fluorescein instead of fixed beads on a slide. On the one hand, this method is cheaper and faster as it does not require a time series to be recorded, also it is insensitive to lateral and axial drift. On the other hand, it has to be prepared fresh each time and is more sensitive to uneven illumination, as all pixels are evaluated for their brightness and contribute to the noise-term. Moreover, the precise positioning of the imaging plane is critical.

For a better reproducibility of the beads method, we advise to exclude or time-crop datasets that display over 5% of drift or bleaching. The overall state of the system should not be under estimated either. It is important to make sure that the general status of the system is comparable between two sets of measurements. For instance, a measurement with a misshaped PSF cannot be compared to one with a good PSF, as the SNR will be expected to be lower. A guide on reproducibility in light microscopy has been recently published with clear guidelines to be followed^27^. Therefore, it is important not to skip this step of the hereby methodology.

While the SBR is a parameter that can be controlled while performing the experiment (by minimizing the image background with better staining protocols, better mounting medium or by applying a digital offset), the SNR is inherent to the imaging process. What parameters influence the SNR scores? First, the strongest influence on the SNR are the number of photons that arrive at a given detector and are converted to photoelectrons. High SNR therefore requires a well aligned light path with minimal light loss (e.g., optimal PSF, proper pinhole alignment, …)^28^ and detectors with a highly effective QE^29^. Second, in PMTs, multiplication noise and dark current contribute to lower SNR but are both functions of gain^29^. A slower scan speed, an optimal gain, a higher laser power and a higher QE will increase the SNR score, and we obtained concordant results with NoiSee when varying those parameters. This is why the SNR results provided by NoiSee need to be seen as scores, obtained with specific acquisition parameters, and not seen as absolute values. A more thorough discussion of which parameters influence SNR and ways to increase SNR on a confocal microscope is presented in the supporting text to this paper (Supplemental Material) and in a publication by Deagle and coworkers ^27^.

What are the minimal SNR and SBR scores needed for a “good” image? When judging image quality, resolution is often the parameter we assess. This can be quantified via a measure of contrast, e.g. the intensity difference between the brightest spots of two objects and the minimum intensity between them. For example, the Rayleigh criterion for resolution corresponds to the distance between two bright objects where 26.4% contrast is achieved^30^. In that regard, GATTAquant nanorulers or Argolight slides, with their associated software are useful tools to address resolution. Contrast is influenced both by SBR and SNR. The influence of SBR is described by the “Weber contrast”, which defines contrast for the human eye as [signal-background]/background^31^. This results in a minimal SBR of 1.264 to fulfil the Rayleigh criterion, i.e. the dimmest signal needs to be 1.264-fold brighter than the background. Considering the signal mean and standard deviation reported in table 1, our measurements reveal that SBR is not a resolution limiting factor for any of the microscopes tested. Assuming a normal distribution of the signal on top of the background, we can estimate the minimal signal to be three standard deviations below the mean (i.e., 0.27% of all values), which is still 20-1000 fold above the minimum contrast criterion. It is noteworthy that albeit the minimal SBR is very small, it still only applies to the human eye. Using a computer, contrasts can always be adjusted for optimal display as shown in Figure 7g-h. Noise on the other hand introduces an uncertainty to the intensity measurement itself which will propagate to the measured contrast, as shown in Figure 7c and 7i. Because of this error, contrast will be reduced in a noisy image, and hence resolution will be reduced as well^1^. Consequently, image resolution and quality degrade continuously with decreasing SNR (Fig4 and 7c/7i). Adjusting a noisy image for optimal display however does not recover any features (Figure7c/7i).

**Figure 7.**
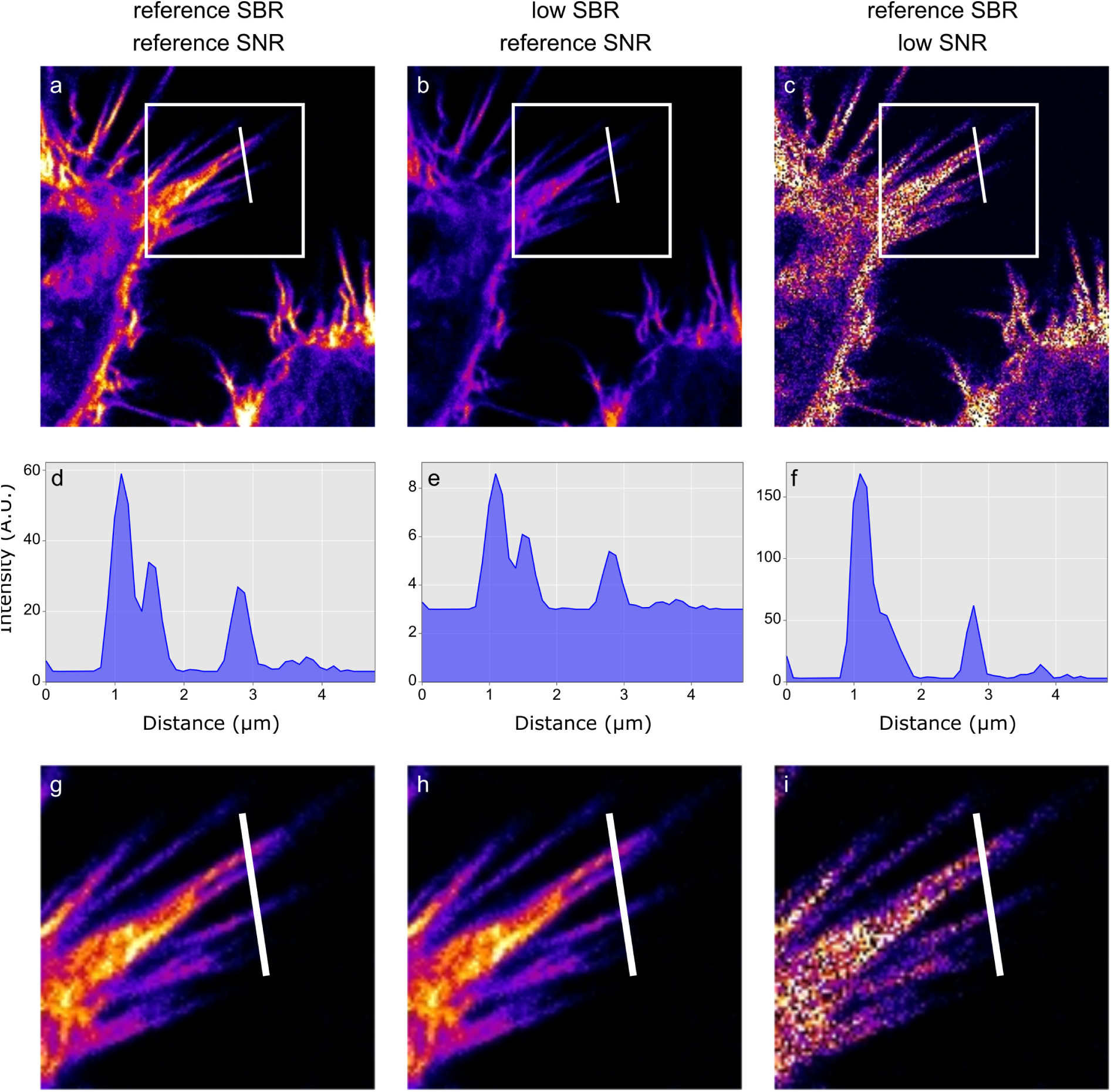
Influence of SNR and SBR on image quality. a. reference image of good SNR and high SBR taken from figure 4b. b. signal in a was lowered 10-fold in Fiji while background and SNR were not lowered. c. same SBR as in a, but a strong Poisson Noise was added to the signal using the RandomJ plugin in Fiji. d. the intensity profile corresponding to the white line in a reveals well-distinguishable features. e. these features are preserved in b despite the low SBR. f. low SNR results in loss of distinguishable features in c. g, h, i. Contrast and brightness were adjusted in Fiji for optimal display in the insets of the corresponding images in a, b, c respectively (white boxes). g. the reference image nicely shows the structure details. h. after brightness and contrast adjustments, the low SBR image looks similar to the reference image in g. i. the brightness and contrast adjustments cannot recover the resolution loss due to the low SNR.

NoiSee was developed for confocal data and validates the known specifications of the different categories of detectors (multialkali PMT, GaAsP PMT and HyD). We were able to show the influence of the sensitivity of specific detectors, and the influence of the instrument noise. The SNR value as calculated using NoiSee is not absolute and is to be understood rather as a score. Similar to the Rose criterion, which states that a SNR of at least 5 is required to distinguish image features by eye^32^, our observations reflect a visible loss of details for systems scoring values below 7 as seen in Figure 4.

NoiSee is the perfect tool to have at hand when routinely checking systems and confocal performances. Here we describe the procedure for confocal microscopes, but the methodology is universal. Resulting scores from confocal and wide-field systems will differ since adjustments in illumination and acquisition settings will be required to allow comparability.

## Material and Methods

### Immunofluorescence staining

HeLa epithelial cells were seeded on 0.17 mm coverslips (#1.5 Zeiss high precision coverslips), fixed for 5 min in 4% PFA and stained with Phalloidin coupled to Alexa 488 (A12379, Molecular Probes, ThermoFisher) for 45 min.

### Beads and Fluorescein slides

A bead standard slide was used for the SNR images (ThermoFisher, T14792). Every microsphere is stained with four different fluorescent dyes [Ex/Em: 365/430 nm (blue), 505/515 nm (green), 560/580 nm (orange), and 660/680 nm (dark red)] that have well-separated excitation and emission peaks. The green 1 µm beads were used for the SNR measurements.

For PSF measurements, beads slides were prepared as follows: 200 nm TetraSpeck microspheres (ThermoFisher, T7280) were diluted 1:6 in ethanol, spread on a #1.5 coverslip and allowed to air-dry. The coverslip was mounted with 5 µl of 100% glycerol on a glass slide, and the edges of the coverslip were sealed with nail polish.

Fluorescein [Ex/Em: 490/514 nm] (46960, Sigma-Aldrich) was dissolved in Milli-Q water to give a 10 mM solution which was stored at 4°C, protected from light. This stock was diluted to 50 µM in water. The solution was added in a depression slide (microscope slide with cavity, #1320000, Marienfeld) and the coverslip was sealed with glue (Twinsil 22, #13001000, Picodent, Wipperfürth, Germany).

### Laser power measurement

To ensure comparability, all measurements were performed at a constant level of illumination (Murray, 1998). Laser power was measured with a Thorlabs power meter (PM200) equipped with a photodiode sensor (S170C, microscope slide format). For each measurement, the microscope system was turned on at least one hour before the measurements were done to allow for thermal stabilization of the system. The surrounding light of the dark room was subtracted by using the “set zero” function of the power meter. The laser power was measured at the image focus plane, when the sensor slide was in contact with the immersion medium (oil). For each AOTF value, the measures were averaged over 50 timepoints (live mode of the power meter). For the laser power measurements to be accurate and comparable between systems, we used the settings described in supplementary table 5. The power used for the study was decided at 1 µW at the image focal plane. Please note that whenever possible, spot scanning was used to measure the most accurate laser power. However, for the LSM800, this functionality is not available, and the correction factor given in supplementary table 5 allows to correct for the blanking time when the measurements are done as specified (256×256 px, maximum speed and maximum zoom).

### Confocal laser scanning microscopy and SNR measurement

Images were collected on a variety of microscopes. In each case, we used a Plan Apo 63x 1.4 numerical aperture (NA) oil objective. Beads were acquired at their centre; the fluorescein solution was imaged at a distance of 10 µm from the coverslip.

*Zeiss LSM700 inverted and upright (point scanning confocals)*. Lens reference 420782-9900-000. Green channel: 488 nm diode laser excitation, variable beam splitter set to 500–700 nm for detection. Detector: multialkali PMT.

*Zeiss LSM800 (inverted point scanning confocal)*. Lens reference 420782-9900-000. Green channel: 488 nm diode laser excitation, variable beam splitter set to 500–700 nm for detection. Detector: GaAsP PMT.

*Leica SP5 (point scanning confocal)*. Lens reference 506192. Green channel: 488 nm diode laser excitation, variable beam splitter set to 500–700 nm for detection. Detectors: multialkali PMT and HyD. In the course of this project, two SP5 microscopes were used, SP5 II and SP5 MP (lens 506350). The SP5 MP was an old system (more than ten years of permanent usage) which only served the purpose of showing low SNR score. The resulting scores are not representative of standard running Leica systems.

*Leica SP8 (point scanning confocal)*. Lens reference 506350. Green channel: 488 nm diode (SP8B) or argon (SP8M) laser excitation, variable beam splitter set to 500–700 nm for detection. Detectors: multialkali PMT and HyD. In the course of this project, two SP8 microscopes were used, SP8B and SP8M. SP8B was used for the fluorescein method, while SP8M was used for the beads method. Please note that SP8M was equipped with white and argon lasers and SMD (Single Molecule Detection) HyDs.

The detection offset was chosen to an insignificant number of zero-value pixels and detector gains were set to optimize the dynamic range while ensuring minimal saturated pixels, i.e. as would have been done for any biological experiment^4^. Detailed settings of the individual instruments are available in supplementary table 1.

### Point spread function acquisition

Bead slides were prepared as described in the “Beads and Fluorescein” section. At the microscope, the following settings were used: 63x oil 1.4 NA objective, frame size 256×256; voxel size 60 nm (x and y) and 200 nm (z); no averaging; pinhole opened to its maximum. 100 z-sections were acquired with the 488 excitation laser. Extra care was taken so the beads were not saturated during acquisition. The resulting PSFs were analysed using the “MIP for PSFs all microscopes” macro^11^.

### SNR calculation in Noisee

Bead method: SNR of an image is calculated as the ratio of the mean pixel value to the standard deviation of the pixel values over a 21 timepoints video of static fluorescent beads. Before SNR calculation we subtract the mean background intensity to account for any offset that may be introduced into the measurement (e.g. detector offset, stray light). Reported is the mean SNR for the highest average intensity value per bead averaged over all beads recorded (as suggested in ^4^).

Fluorescein method: After subtraction of the mean intensity in the dark image from the Fluorescein image, the SNR is calculated as the ratio of the mean intensity divided by its standard deviation of all pixels in the background-subtracted Fluorescein image.

### Data analysis

The data was plotted and statistical analysis was performed in Python using Welch’s *t*-test method provided by SciPy^33^.

## Supporting information

## Acknowledgments

We thank the Zeiss team for their advices on this project, especially Christine Strasser, Ingo Bartholomaeus and Andreas Seher. We also thank Kurt Thorn and Nico Stuurman (UCSF) for their original method to perform SNR measurements as well as Adam Mazur for his valuable input on data analysis. No beads were harmed during the process of the acquisition.

## Author contributions statement

AF, KDS and OB conceived and designed the experiments, AF performed the experiments, KDS created the analysis tool, NE revised the analysis tool. AF, KDS, NE, WH and OB analysed and interpreted the results. AF, KDS and OB wrote the manuscript.

## Competing interests

The authors declare no competing interests.

## Notes

#### Summary of Updates

The title has been changed. Text and figures have been changed as well.

