## Supplementary material for "Using the NoiSee workflow to measure signal-to-noise ratios of confocal microscopes"

### Supplementary figures

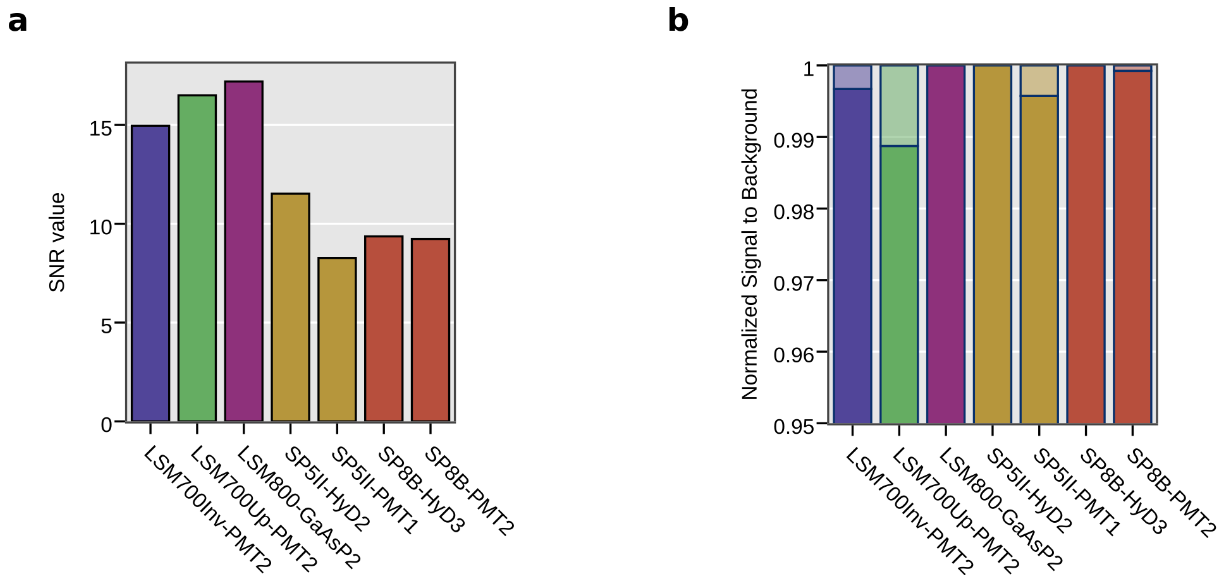


**Supplementary figure 1** – Results obtained by the Fluorescein method. **a.** NoiSee SNR scores for 7 detectors across the 5 different tested systems. **b.** NoiSee SBR scores for the same detectors and systems. Solid colour denotes the mean signal while transparent colour denotes the mean background measured on a fluorescein image.

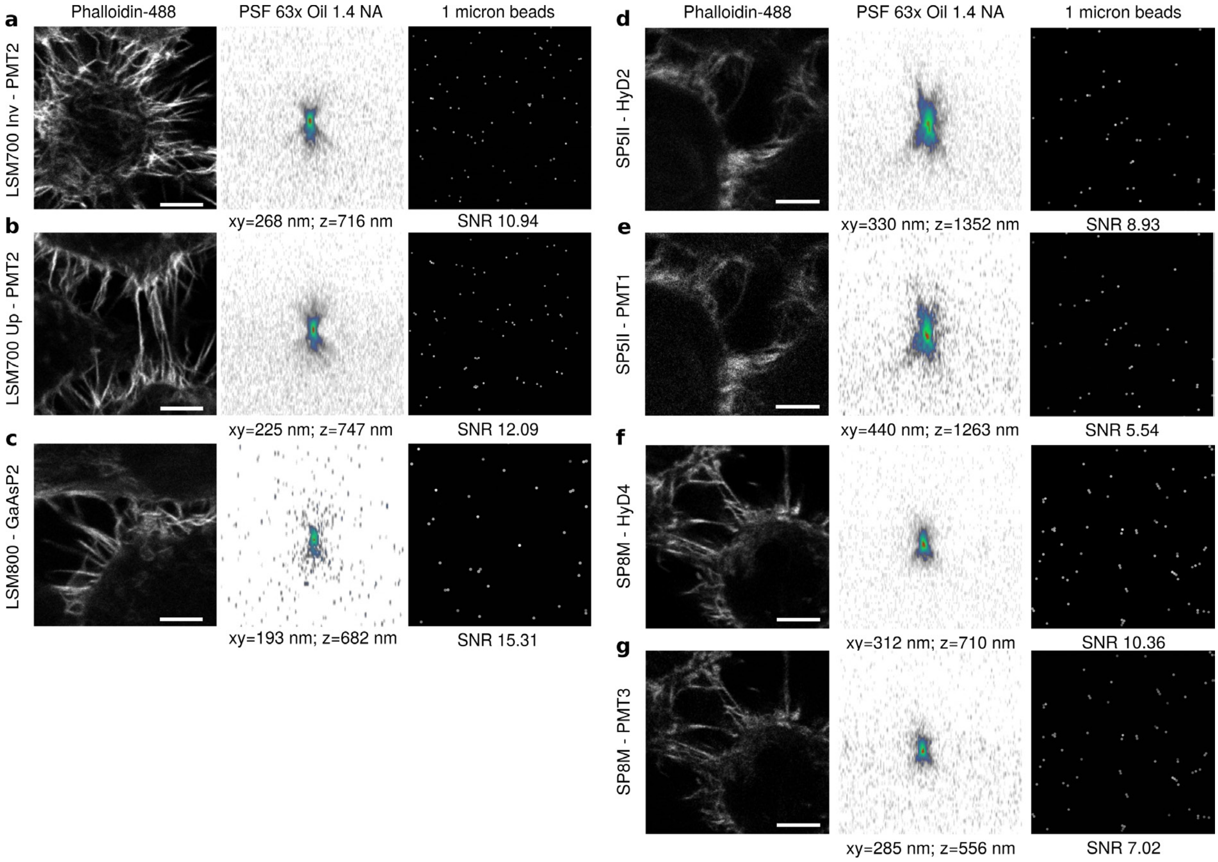


**Supplementary figure 2** - ID-cards of the tested detectors for the beads method. For each detector, the left panel shows a zoom from the “general check-up” image (cells stained with phalloidin-488). The central panel depicts the PSF and its associated lateral and axial full width at half maximum values. The right panel shows the beads images and associated NoiSee SNR scores. **a/b.** Standard Zeiss PMT detectors. **c.** Zeiss GaAsP. **d/f.** Leica HyD detectors. **e/g.** Standard Leica detectors.

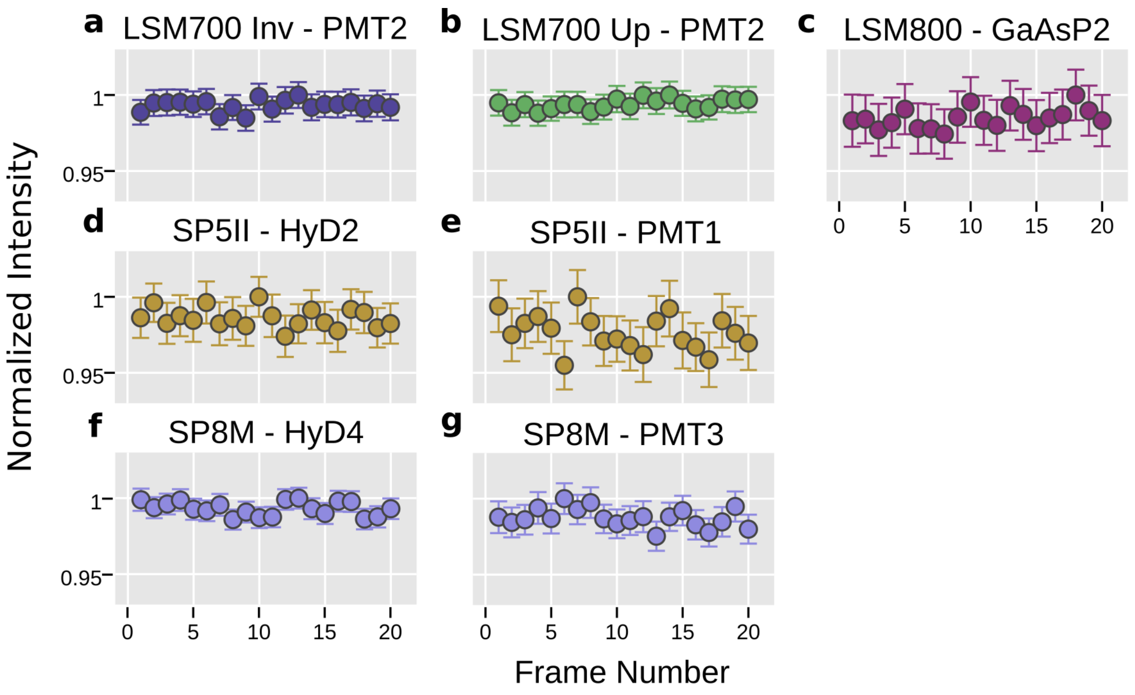


**Supplementary figure 3** – Normalized mean intensity across all beads within one frame as measured by NoiSee for all instruments and detectors analysed. The intensity fluctuations generally stay within the measurement error, proving that beads did not bleach during acquisition time. The first datapoint was excluded in order to avoid influence of non-equilibrium effects[1]. Error bars represent the standard error of the mean.

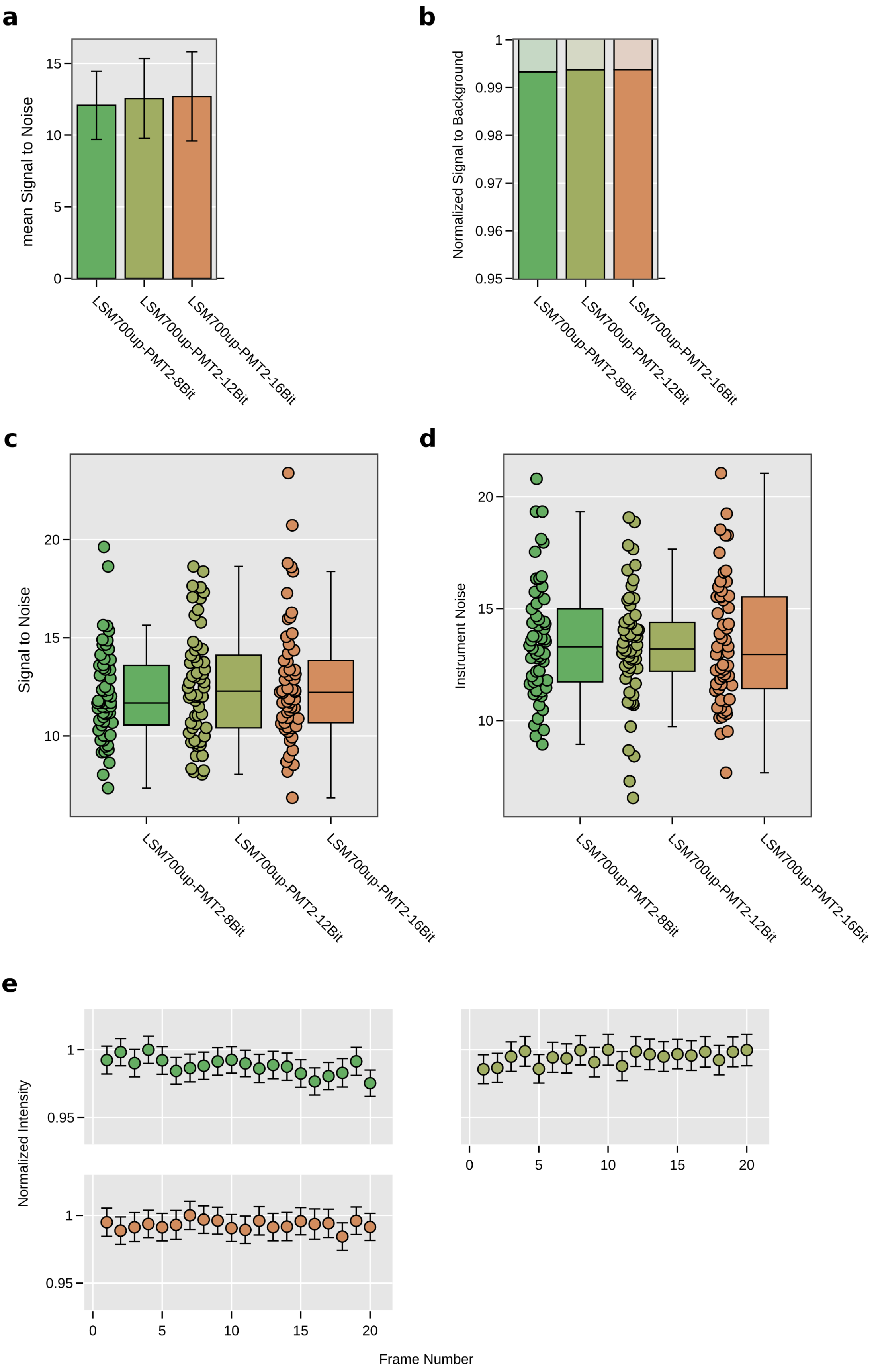
a
b
c
d
e
mean Signal to Noise
Normalized Signal to Background
Signal to Noise
Instrument Noise
Normalized Intensity
Frame Number
LSM700up-PMT2-8Bit
LSM700up-PMT2-12Bit
LSM700up-PMT2-16Bit

**Supplementary figure 4** – Repeated measurements of beads acquired at various bit depths in the same field of view on the LSM700up. **a.** Bar charts of mean SNR score at 8-, 12-, and 16-bit (green, olive, orange. Welch's t-test for independent samples revealed no significant differences [p(8-bit vs. 12-bit)=0.33, p(8-bit vs. 16-bit)=0.23, p(12-bit vs. 16-bit)=0.79]. SBR of the measurements corresponding to the data presented in **a**. The distribution of individual values is summarize in the boxplots presented in **c** (SNR score) and **d** (noise), respectively. **e**. Normalized mean intensity across all beads measured reveals that intensity fluctuations stay within the measurement error, indicating no bleaching during acquisition. Error bars represent the standard deviation (**a**) or the standard error of the mean (**e**).

### Supplementary tables

**Supplementary table 1** - Settings used at the different microscopes to ensure comparability of the beads results

| | LSM700 Inv (PMT2) | LSM700 Up (PMT2) | LSM800 (GaAsP2) | SP5 II (PMT1 / HyD2) | SP8 M (PMT3 / HyD4) |
|---|---|---|---|---|---|
| Frame size; zoom | 1024; x1 | 1024; x1 | 670; x1.5 | 800; x3.1 | 800; x2.3 |
| Resulting xy pixel size (nm) | 99.2 | 99.2 | 100.9 | 99.3 | 100.4 |
| Scanner speed | 7 | 7 | 7 | 200 Hz | 200 Hz |
| Pixel Dwell Time (µsec) | 1.576 | 1.576 | 1.58 | 1.550 | 1.550 |
| Pinhole size (µm; given by software) | 40.86 | 40.84 | 40.92 | 85.66 | 85.60 |
| Backprojected PH radius (nm; SVI calculator) | 239.08 | 238.96 | 239.43 | 255.71 | 255.53 |
| Detection range (nm) | 500-700 | 500-700 | 500-700 | 500-700 | 500-700 |
| Laser power / HyD Gain | ********** | ********** | ********** | 2% / 35% | 0.95% / 20% |
| Laser power / GaAsP Gain / offset | ********** | ********** | 0.04% / 540 / 1 | ********** | ********** |
| Laser power / PMT Gain / offset | 0.2% / 600 / 2 | 0.2% / 650 / 2 | ********** | 2% / 750 / -0.33 | 0.95% / 700 / -0.09 |

**Supplementary table 2** - Settings used at the different microscopes to ensure comparability of the Fluorescein results

| | LSM700 Inv (PMT2) | LSM700 Up (PMT2) | LSM800 (GaAsP2) | SP5 II (PMT1 / HyD2) | SP8 B (PMT2 / HyD3) |
|---|---|---|---|---|---|
| Frame size; zoom | 512; x2 | 512; x2 | 670; x1.5 | 800; x3.1 | 800; x2.3 |
| Resulting xy pixel size (nm) | 99.2 | 99.2 | 100.9 | 99.3 | 99.3 |
| Scanner speed | 7 | 7 | 7 | 200 Hz | 200 Hz |
| Pixel Dwell Time (µsec) | 1.576 | 1.576 | 1.58 | 1.550 | 1.550 |
| Pinhole size | 40.86 | 40.84 | 40.92 | 85.66 | 85.66 |

| | | | | | |
|---|---|---|---|---|---|
| (µm; given by software) | | | | | |
| Backprojected PH radius (nm; SVI calculator) | 239.08 | 238.96 | 239.43 | 255.71 | 255.71 |
| Detection range (nm) | 500-700 | 500-700 | 500-700 | 500-700 | 500-700 |
| Laser power / HyD Gain | ********** | ********** | ********** | 2% / 15% | 0.08% / 10% |
| Laser power / GaAsP Gain / offset | ********** | ********** | 0.03% / 500 / 0 | ********** | ********** |
| Laser power / PMT Gain / offset | 0.2% / 550 / 2 | 0.2% / 600 / 2 | ********** | 2% / 675 / -0.33 | 0.08% / 600 / -0.13 |

**Supplementary table 3** - Results at the different microscopes using the Fluorescein method

| | LSM700 Inv (PMT2) | LSM700 Up (PMT2) | LSM800 (GaAsP2) | SP5 II (HyD2) | SP5 II (PMT1) | SP8 B (HyD3) | SP8 B (PMT2) |
|---|---|---|---|---|---|---|---|
| Mean signal | 47848.01 | 193.08 | 180.11 | 156.04 | 174.78 | 53.13 | 79.22 |
| Noise | 3197.57 | 11.7 | 10.47 | 13.55 | 21.13 | 5.68 | 8.58 |
| SNR | 14.96 | 16.5 | 17.2 | 11.52 | 8.27 | 9.36 | 9.23 |
| Background mean | 158.71 | 2.2 | 0.0002429 | 0.000019 | 0.75 | 2.316e-05 | 0.06 |
| SBR | 301.47 | 87.75 | 741436.58 | 8322148.5 | 233.4 | 2293901.33 | 1247.97 |
| Background standard deviation | 173.84 | 0.52 | 0.02 | 0.01 | 0.47 | 0.0 | 0.25 |

**Supplementary table 4** – NoiSee results of repeated bead measurements on two different microscopes.

| | LSM700 Inv (PMT2) R1 | LSM700 Inv (PMT2) R2 | LSM700 Up (PMT2) R1 | LSM700 Up (PMT2) R2 |
|---|---|---|---|---|
| SNR mean | 10.94 | 9.5 | 12.09 | 12.03 |
| SNR standard deviation | 2.55 | 1.7 | 2.03 | 2.45 |
| SBR mean | 90.68 | 75.86 | 86.26 | 81.51 |
| Signal mean | 148.89 | 127.08 | 167.18 | 158.0 |
| Signal standard deviation | 29.39 | 24.4 | 29.48 | 30.1 |
| Background mean | 1.64 | 1.68 | 1.94 | 1.94 |
| Background standard deviation | 0.55 | 0.51 | 0.42 | 0.39 |
| n beads | 66 | 53 | 56 | 52 |

**Supplementary table 5** – Description of the methodology used to measure laser power at the image focal plane

| Confocal system (Manufacturer) | Methodology | Correction factor |
|---|---|---|
| SP5 (Leica) | Bleachpoint, 1400 Hz | - |
| SP8 (Leica) | Bleachpoint, 1800 Hz | - |
| LSM700 (Zeiss) | Spot scanning | - |
| LSM800 (Zeiss) | Maximum speed scanning: 256x256, Max Speed (16), Zoom max | Divide the measured power by 0.836 to correct for the blanking time |

*Supplementary Table 6: Results at the different bit depths using the Beads method*

| | LSM700 Up (PMT2-8bit) | LSM700 Up (PMT2-12bit) | LSM700 Up (PMT2-16bit) |
|---|---|---|---|
| SNR mean | 12.079 | 12.5538 | 12.6997 |
| SNR standard deviation | 2.3801 | 2.7847 | 3.1144 |
| SBR mean | 160.931 | 158.602 | 159.937 |
| Signal mean | 148.1301 | 162.6552 | 165.2414 |
| Signal standard deviation | 30.1341 | 32.0551 | 30.9007 |
| Background mean | 1.0864 | 1.0256 | 1.0332 |
| Background standard deviation | 0.3564 | 0.1989 | 0.2141 |
| n beads | 58.0 | 58.0 | 58.0 |

### NoiSee installation and user guide

#### Installation

- Download and install Fiji as described at https://imagej.net/Fiji/Downloads
- NoiSee is available from its ImageJ update site and can be installed following the official guide at https://imagej.net/Following_an_update_site

#### User guide – Bead method

- Start Fiji
- Type “NoiSee” in the quick search bar, select “Bead Analysis” and click “Run”

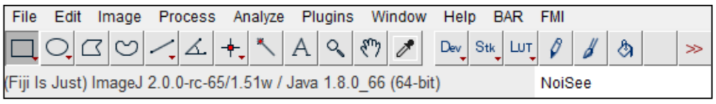


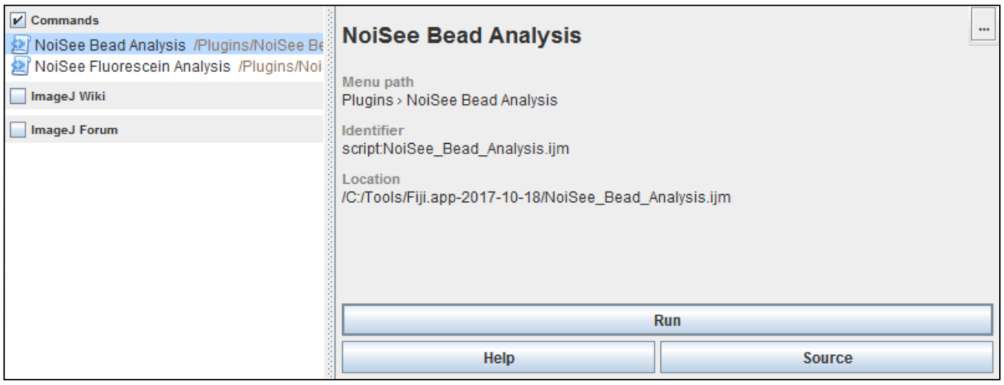


In the analysis dialog,

- Specify the path to the time lapse bead image.
  - *Note: NoiSee uses Bio-Formats*[2] *to open images and hence can be used directly with many native file formats like Zeiss *.lsm or Leica *.lif.*
  - All result files generated by NoiSee will be placed in a new sub-folder located in the same directory as the input image.
- Specify the diameter of the beads in pixels.
- NoiSee uses the “Find Maxima” function to automatically detect beads above a certain intensity threshold that needs to be roughly estimated. Starting values for this estimate depending on the detector type used, e.g. 50 for PMTs, 10 for Hybrid detectors in photon counting mode or 500 for camera based systems.
  - If not all beads are identified the estimate should be lower.
  - If peaks other than beads are identified the estimate should be higher.
- *Note: While NoiSee tries to exclude hot pixels and overexposed beads, these would ideally be already avoided during imaging.*
- Choose whether you wish to create kymographs. We recommend to do so as they are useful to inspect sample drift and bleaching.
- NoiSee can further save additional data (individual measurements per bead) as txt-files.

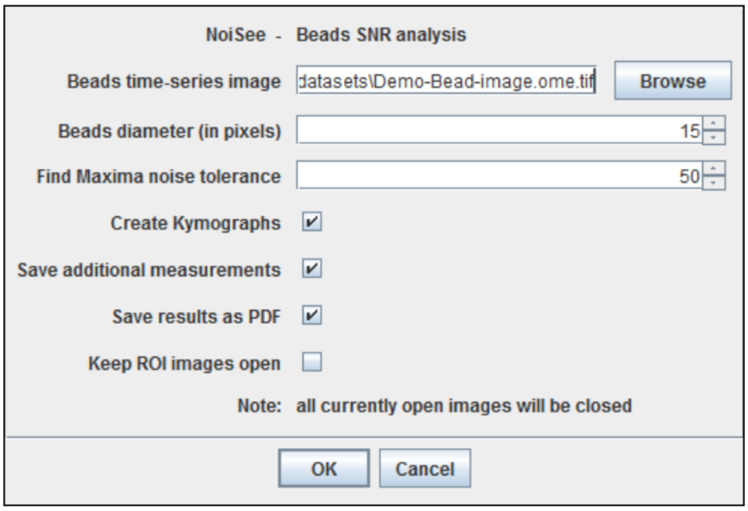


- All images and plots created by NoiSee can optionally be saved in a PDF report.
- ROIs used for measurements are available from the ROI manger and can additionally be visualized as extra images.
- Press “OK” and wait for NoiSee to finish its calculations. The final output which will be arranged as presented in Figure 3 of the main manuscript.
- A summary of all NoiSee results is automatically saved as a txt-file.

#### User guide – Fluorescein method

- Start NoiSee as above but select “Fluorescein Analysis” instead of “Bead Analysis”.
- Specify the path to the dark image and fluorescein image.
    - All result files generated by NoiSee will be placed in a new sub-folder located in the same directory as the input image.

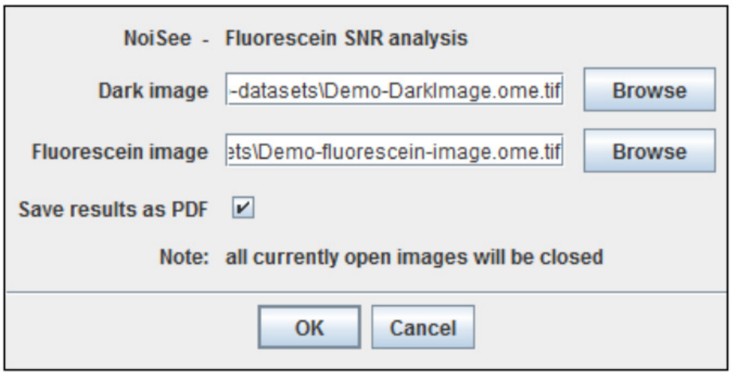


- Again all images and plots created by NoiSee can optionally be saved in a PDF report.
- Press “OK” and wait for NoiSee to finish its calculations as presented in the screenshot below.

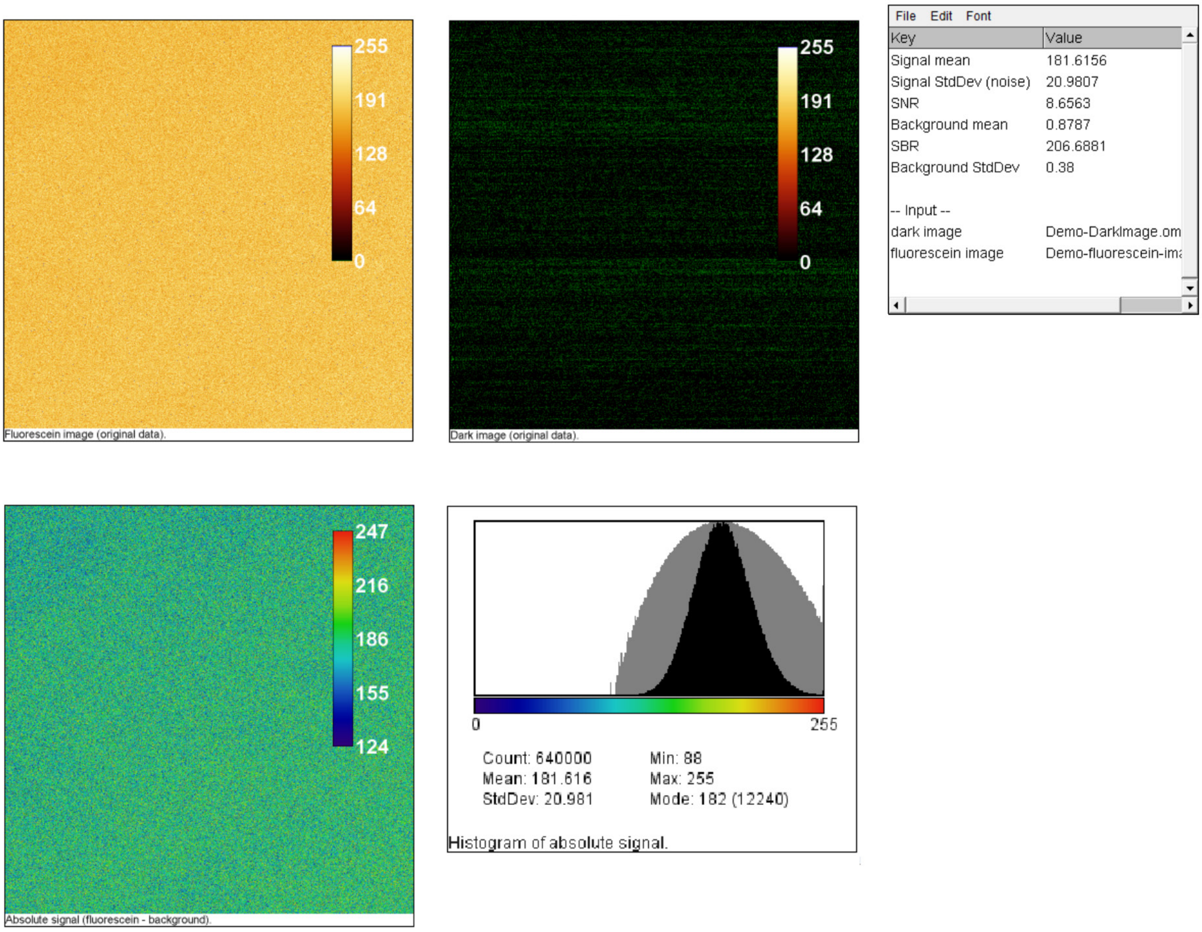


Screenshot of NoiSee output. From left to right and top to bottom: raw Fluorescein image colouring zero values in green (clipping), and overexposed pixels in blue; raw dark image viewed with the same lookup table; Results table presenting a summary of the measurements; Fluorescein image subtracted by the mean intensity of the dark image; Histogram of the background-corrected Fluorescein image.

### Supporting Text

In theory the SNR from a detector can be described as

$$SNR = \frac{N * QE}{\sqrt{N * QE + D}}, \qquad D = M + d + E^2 \qquad \textit{Eq. 1}$$

where N*QE denotes the number of incident photons N converted to photoelectrons (i.e. the signal) and D the cumulated detector noise, i.e. multiplication noise (M), dark current (d) and electronic noise (E)[3,4]. As the photon shot noise follows a Poisson-distribution, it is described by the square root of the signal.

The strongest influence on the SNR is the number of photons N that arrive at the detector, which rises in response to increasing laser power. While this number may vary between systems due to their intrinsic light path which usually is not alterable by the user, drastic decay of the SNR seen from repeated measurements of the same detector using constant laser power indicates that one or several elements in the beam path are out of alignment, e.g. the pinhole or laser itself.

Higher SNR also results from lowering the dark current and multiplication noise when PMTs are used. While higher gains generally lower the multiplication noise[5], they also enhance the dark current due to additional generation of thermal electrons. Hence, each detector has an optimal gain setting. This optimal setting can be identified when plotting the dark current as a function of gain. Beyond a certain voltage, the dark current is rising nonlinearly and should therefore not be set higher than this value. Hence, further improvements are possible when cooled detectors are used.

A common strategy to increase SNR in confocal laser scanning microscopy is to use signal averaging or accumulation. This may however contribute to additional image degradation that might be interpreted as noise but in fact results from a sample drift in the xy-plane. Drift in this case will lead to an apparent blurring or smearing of the imaged structures that will be more pronounced with increasing acquisition time. It is possible to check if extensive drift is present by inspecting the kymographs and the plot of mean bead intensity over time as created by NoiSee. Note that z-drift is indistinguishable from bleaching on short timescales.
